# MAMGL: A memory-augmented meta-graph learning framework for adolescent major depression disorder diagnosis

**DOI:** 10.64898/2026.03.27.714607

**Authors:** Xiaobo Liu, Xin Wen, Liang He, Xiaoqiang Liu, Yujun Gao, Xianwei Guo

## Abstract

**Background:** Adolescent major depressive disorder (AMDD) is a prevalent and heterogeneous psychiatric condition that emerges during a critical period of brain development. Neuroimaging-based biomarkers derived from resting-state functional magnetic resonance imaging (rs-fMRI) hold promise for objective diagnosis; however, pronounced inter-individual variability and limited sample sizes pose major challenges for robust model development.

**Methods:** We propose a memory-augmented Meta-Graph Convolutional Network (BrainMetaGCN) to classify AMDD using rs-fMRI functional connectivity. Individual functional connectivity matrices were constructed by parcellating rs-fMRI time series into cortical regions of interest and computing pairwise correlations. A meta-graph generator dynamically learned subject-specific graph structures, which were processed by lightweight graph convolutional layers. A memory neural network was incorporated to encode population-level prototypical connectivity patterns and generate individualized representations via attention-based retrieval. Model performance was evaluated across multiple independent datasets and compared with state-of-the-art deep learning approaches. Additionally, network interpretability was examined through cortical hierarchy analysis and functional enrichment of discriminative network components.

**Results:** The proposed BrainMetaGCN consistently outperformed baseline models, including convolutional and transformer-based approaches, achieving higher accuracy, area under the receiver operating characteristic curve, sensitivity, and specificity. Memory-module–derived functional networks exhibited clear modular organization and showed a significant positive correlation with cortical functional hierarchy, supporting their neurobiological validity. Functional enrichment analyses implicated synaptic transmission, axon guidance, receptor tyrosine kinase signaling, and immune-related pathways, suggesting neurodevelopmental and neuroimmune mechanisms underlying AMDD. Ablation analyses confirmed that memory augmentation and dynamic meta-graph construction were critical for robust performance under small-sample conditions.

**Conclusions:** This study introduces a robust and interpretable memory-augmented graph learning framework for AMDD classification. By effectively balancing individual specificity and population-level generalization, BrainMetaGCN advances neuroimaging-based precision diagnosis and provides new insights into the neural and biological mechanisms of adolescent depression.

## Introduction

Adolescent major depressive disorder (AMDD) is a common psychiatric condition characterized by persistent low mood, anhedonia, and cognitive–behavioral dysfunction, with its incidence steadily rising worldwide (*1*, *2*). Adolescence is a critical period of neurodevelopment, during which the prefrontal cortex remains functionally immature, constraining emotion regulation and impulse control and thereby increasing susceptibility to environmental stressors. Concurrent hormonal and neurotransmitter fluctuations, together with academic, interpersonal, and familial pressures, further amplify the risk of depressive symptoms (*3*). Clinically, diagnosis still relies predominantly on symptom rating scales (e.g., PHQ-9, CDI) and structured or semi-structured interviews. However, marked heterogeneity in clinical presentation—from typical low mood to prominent irritability, attentional difficulties, or somatic complaints—combined with variability in adolescents’ capacity for self-expression, frequently leads to misdiagnosis and underdiagnosis (*4*).

In contrast, neuroimaging-based objective approaches, such as functional magnetic resonance imaging (fMRI), offer a noninvasive and temporally sensitive means of capturing brain activity, providing an important avenue for elucidating the neural mechanisms underlying depression (*5*). fMRI measures blood oxygen level–dependent (BOLD) signal fluctuations associated with neuronal activity and thus indirectly reflects the dynamic organization of large-scale functional brain networks over time (*6*, *7*). Prior studies have documented significant alterations in functional connectivity within the default mode network (DMN), executive control network (ECN), and other key brain regions in adolescents with depression (*6*). These macroscopic network abnormalities may be closely linked to microscopic processes, including neurotransmitter receptor distributions and gene-regulatory mechanisms (*6*). Nonetheless, the pronounced inter-individual variability in both clinical symptomatology and functional connectivity profiles in AMDD presents a major methodological challenge: how to robustly extract and preserve informative connectivity features across individuals.

Recent progress in deep learning—particularly graph neural network–based approaches—has created new opportunities for the analysis of brain functional networks (*8*). Graph convolutional networks (GCNs) can effectively exploit interregional connectivity to model resting-state fMRI (rs-fMRI) data and have achieved promising performance in the diagnostic classification of several psychiatric disorders, including MDD. By constructing functional connectivity graphs and applying GCNs to derive graph-level features, previous studies have reported high accuracy in distinguishing patients from healthy controls (*8*). However, these methods often require large sample sizes and assume relatively stable prior graph structures, which constrains their generalizability and robustness in AMDD research, where datasets are typically small and inter-individual variability is substantial (*9*). To overcome these limitations, memory neural networks (MNNs) have emerged as a promising framework for individualized modeling(*10*, *11*). Without relying on prior structural information, MNNs can compress population-level connectivity patterns into a set of “prototypical templates” and represent each individual as a unique combination of these templates, thereby retaining individual-specific characteristics while preserving population-level generalizability. Moreover, by integrating historical state information, MNNs can generate connectivity representations that reflect different stages of illness progression. When coupled with low-rank factorization, this approach effectively reduces model complexity, helping to mitigate the high noise and small-sample constraints of rs-fMRI data and thereby improving diagnostic stability and robustness in previously unseen individuals (*12*, *13*).

Against this background, the present study seeks to integrate lightweight deep learning models with functional brain network analysis to develop an efficient, interpretable, and robust classification framework for AMDD. Specifically, we propose an enhanced lightweight GCN architecture augmented with a memory network mechanism to extract prototypical functional connectivity patterns from rs-fMRI data in adolescents with depression. The learned network patterns are subsequently mapped onto established brain parcellation atlases to provide biological and neuroanatomical interpretation of the discriminative features. Finally, to rigorously evaluate the robustness and generalizability of the proposed framework, we conduct systematic experiments under diverse hyperparameter settings, classifier configurations, and across multiple independent datasets. This work not only offers a novel methodological avenue for the precise diagnosis of AMDD, but also contributes new insights into its underlying neural mechanisms.

## Results

### Overview of the proposed memory-augmented Meta-GCN framework

We developed a memory-augmented Meta-GCN framework to address the high inter-individual variability and small-sample challenges inherent in adolescent major depressive disorder (AMDD) neuroimaging studies (Figure 1). By dynamically constructing subject-specific functional graphs and integrating a memory module to encode population-level prototypical connectivity patterns, the proposed model aims to learn robust yet individualized representations from resting-state fMRI data. This lightweight architecture enables effective extraction of discriminative functional connectivity features while maintaining interpretability and generalizability.

**Figure 1.**
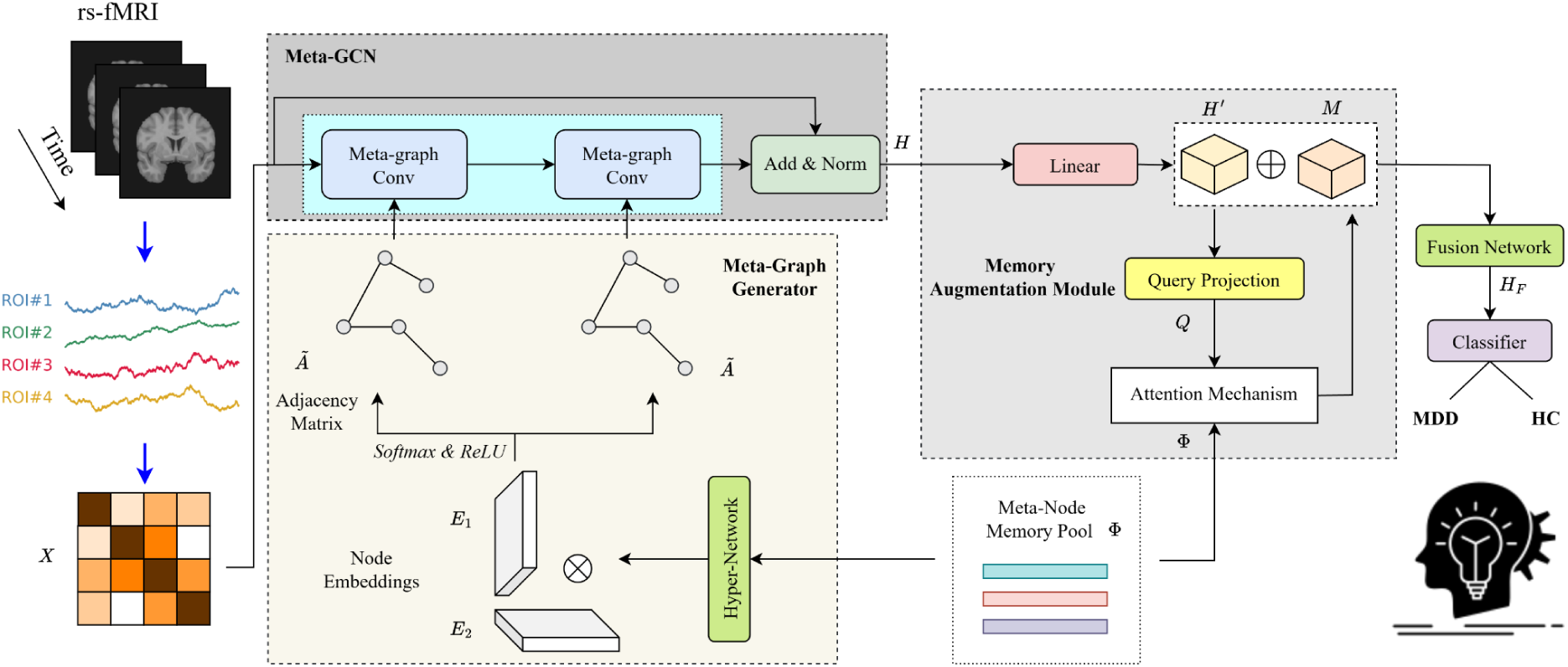
Overall architecture of the proposed memory-augmented Meta-GCN framework for AMDD classification. Schematic overview of the proposed memory-module–based Meta-GCN framework. Resting-state fMRI time series are first parcellated into cortical regions of interest (ROIs) and transformed into individual functional connectivity matrices. A meta-graph generator dynamically constructs subject-specific adjacency matrices based on learned node embeddings, which are subsequently processed by stacked meta-graph convolutional layers with residual normalization. To enhance individualized representation learning under high inter-subject variability, a memory augmentation module is introduced to encode population-level prototypical connectivity patterns and retrieve subject-adaptive representations via an attention mechanism. The fused representations are finally fed into a classifier to discriminate adolescents with major depressive disorder (MDD) from healthy controls (HC).

This study included two independent datasets (Supplementary Table 1). The exploratory dataset comprised 302 adolescents with major depressive disorder (MDD; mean age = 15.40 ± 2.09 years; 134 females) and 207 healthy controls (HC; mean age = 15.53 ± 1.98 years; 82 females). All participants were right-handed, of Han Chinese ethnicity, and had no history of other psychiatric disorders or major physical illnesses. Resting-state fMRI data from all participants underwent identical quality control procedures, including the exclusion of scans with excessive head motion or imaging artifacts. Subsequently, individual fMRI time series were parcellated into 400 cortical regions using the Schaefer atlas, and Pearson correlation coefficients were computed between all pairs of regions to generate subject-specific functional connectivity matrices. Group-level functional connectivity matrices were then obtained by averaging the individual connectivity matrices across participants.

### Superior classification performance compared with state-of-the-art methods

We first evaluated the classification performance of the proposed BrainMetaGCN model against several representative baseline approaches, including BrainNetCNN, Vanilla Transformer (VanillaTF), BrainNetTF, and BrainGCN (Figure 2). Across all evaluation metrics—accuracy (ACC), area under the receiver operating characteristic curve (AUC), sensitivity (SEN), and specificity (SPEC)—BrainMetaGCN consistently outperformed competing methods.

**Figure 2.**
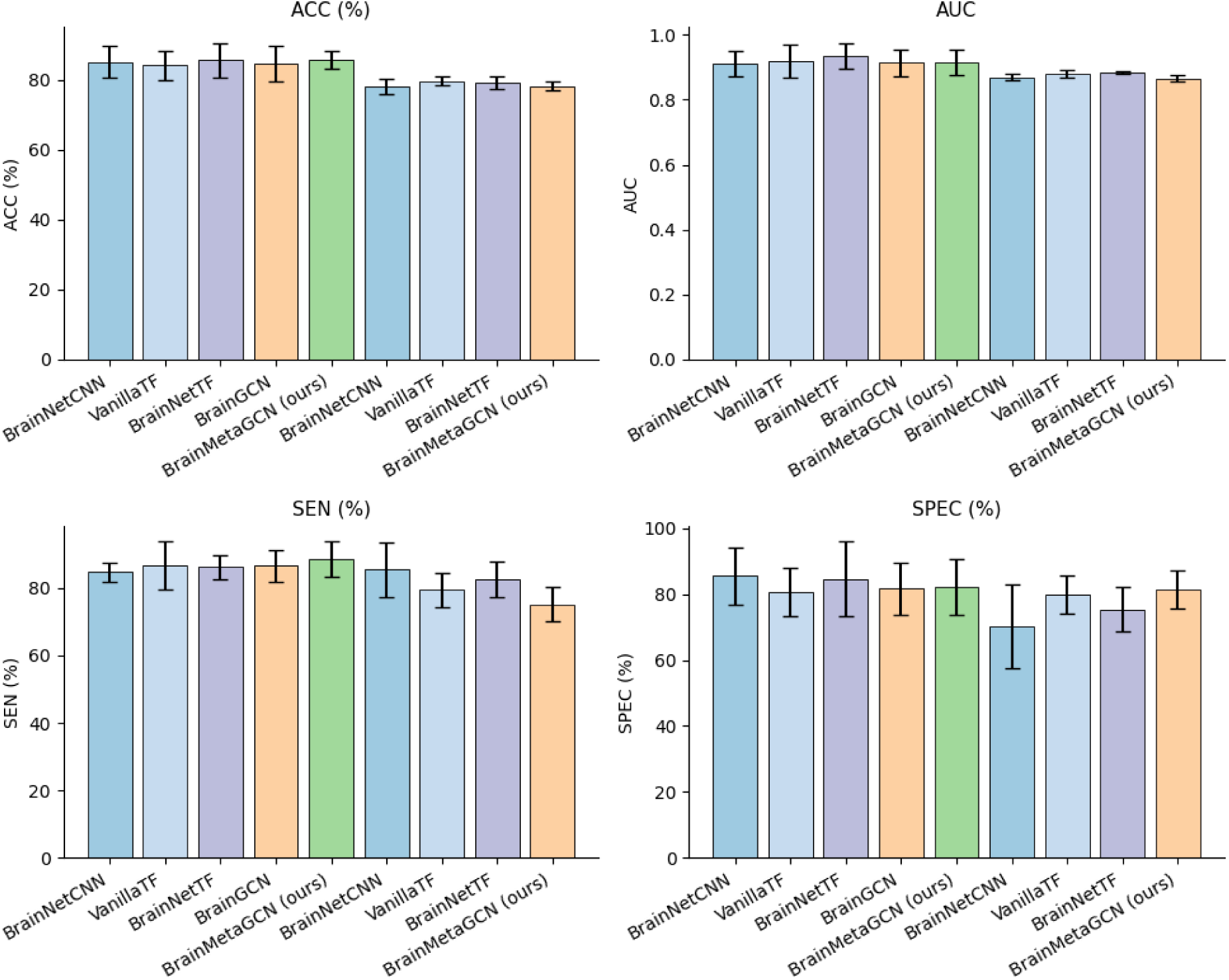
Classification performance comparison with state-of-the-art methods. Performance comparison between the proposed BrainMetaGCN model and baseline approaches, including BrainNetCNN, VanillaTF, BrainNetTF, and BrainGCN, evaluated using accuracy (ACC), area under the ROC curve (AUC), sensitivity (SEN), and specificity (SPEC). Bar plots report mean performance across cross-validation folds, with error bars indicating standard deviation. The proposed BrainMetaGCN consistently outperforms competing methods across all evaluation metrics, demonstrating improved robustness and discriminative capability for adolescent MDD classification.

Specifically, the proposed model achieved the highest mean ACC and AUC, indicating improved overall discriminative ability and robustness. Notably, BrainMetaGCN also demonstrated superior sensitivity, suggesting enhanced capability to correctly identify adolescents with MDD, while maintaining competitive specificity. These results indicate that integrating meta-graph construction with memory-based representation learning substantially improves diagnostic performance under conditions of pronounced heterogeneity and limited sample size.

### Memory-module–derived functional networks capture biologically meaningful organization

To examine the internal representations learned by the proposed framework, we visualized the memory-module–based functional connectivity patterns (Figure 3a). Compared with raw correlation matrices, the reconstructed connectivity matrices exhibited clearer structure and reduced noise, reflecting coherent inter-regional interactions. Graph visualizations further revealed modular organization within the learned networks, suggesting that the memory module effectively distilled stable population-level connectivity motifs while preserving individual-specific variations.

**Figure 3.**
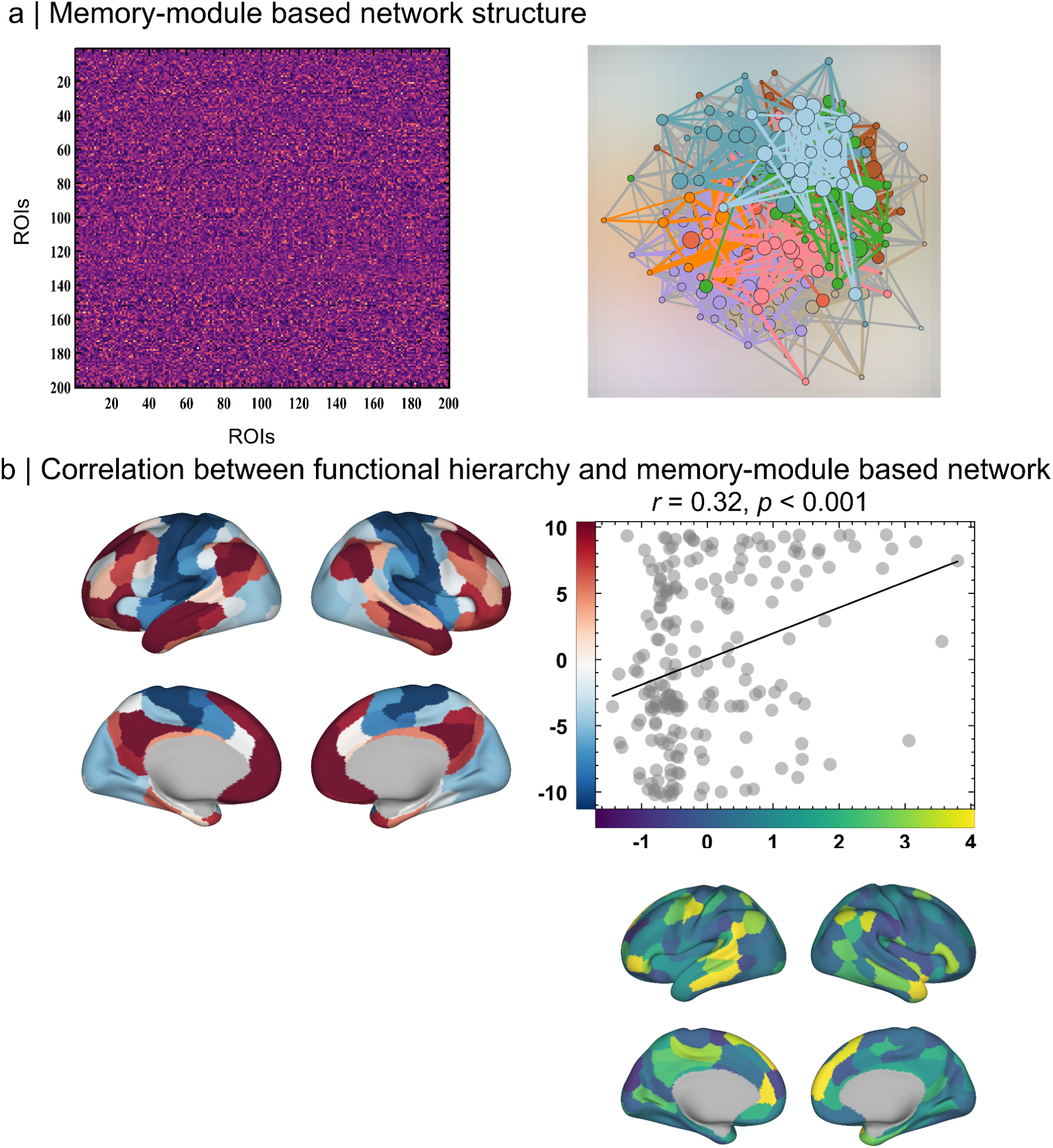
Visualization of memory-module–based functional network representations. **(a)** Left: Representative functional connectivity matrix reconstructed from memory-module–based representations, illustrating denoised and structured inter-regional connectivity patterns. Right: Graph visualization of the corresponding memory-informed functional brain network, where nodes represent ROIs and edges reflect learned functional associations. Distinct colors denote network-level modular organization derived from the memory module. **(b)** Relationship between cortical functional hierarchy and memory-module–derived network representations. Surface maps illustrate regional hierarchy scores and memory-based network weights projected onto the cortical surface. Scatter plot shows a significant positive correlation between functional hierarchy and memory-module representations (r = 0.32, p < 0.001), indicating that the learned memory templates capture biologically meaningful hierarchical organization of brain networks.

To assess the biological relevance of these representations, we examined their relationship with cortical functional hierarchy (Figure 3b). A significant positive correlation was observed between memory-module–derived network weights and established functional hierarchy measures (r = 0.32, p < 0.001). Regions with higher hierarchical positioning tended to exhibit stronger contributions within the memory-informed networks. This finding suggests that the learned connectivity templates are not arbitrary but align with known principles of large-scale cortical organization, supporting the neurobiological plausibility of the proposed approach.

### Functional enrichment analysis reveals neurodevelopmental and synaptic mechanisms

We further investigated the biological significance of the discriminative network components identified by BrainMetaGCN through functional enrichment analysis (Figure 4). The resulting enrichment network revealed clusters of biological processes and signaling pathways highly relevant to the pathophysiology of depression.

**Figure 4.**
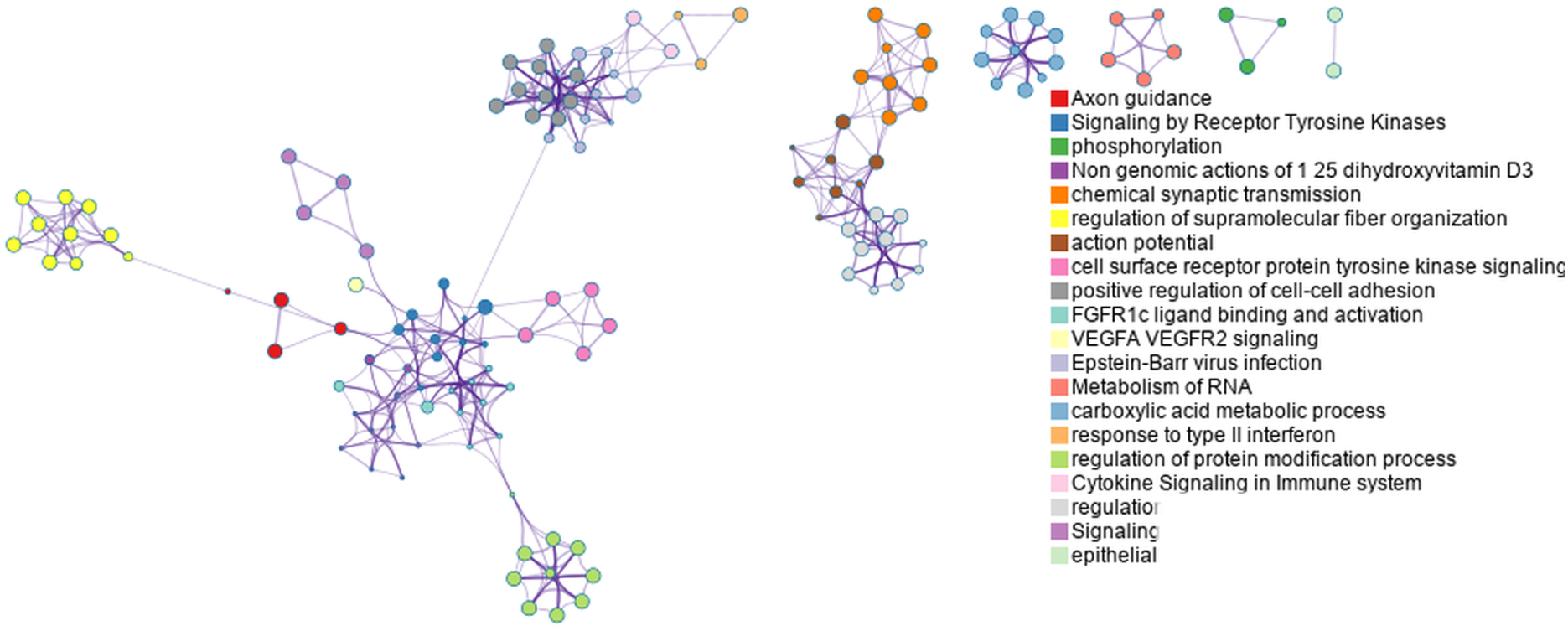
Functional enrichment analysis of discriminative network components. Gene ontology and pathway enrichment network derived from brain regions and connections identified as highly discriminative by the proposed model. Nodes represent enriched biological processes or signaling pathways, and edges indicate functional similarity or shared gene involvement. Prominent clusters are associated with synaptic transmission, axon guidance, receptor tyrosine kinase signaling, immune-related cytokine signaling, and neurodevelopmental processes, suggesting potential molecular and cellular mechanisms underlying adolescent major depressive disorder.

Prominent enriched terms included synaptic transmission, axon guidance, receptor tyrosine kinase signaling, and regulation of cell–cell adhesion, highlighting alterations in neural communication and network development. In addition, pathways related to immune and cytokine signaling were identified, consistent with growing evidence implicating neuroimmune interactions in depressive disorders. Collectively, these findings suggest that the functional connectivity patterns captured by the proposed model may reflect underlying molecular and cellular mechanisms associated with neurodevelopmental dysregulation in AMDD.

### Ablation and hyperparameter analyses demonstrate robustness and stability

To evaluate the robustness of the proposed framework and the contribution of key model components, we conducted systematic ablation and hyperparameter sensitivity analyses (Figure 5). Performance comparisons across different node embedding dimensions and numbers of meta-graph supports demonstrated that BrainMetaGCN maintained stable and high classification performance across a broad range of settings.

**Figure 5.**
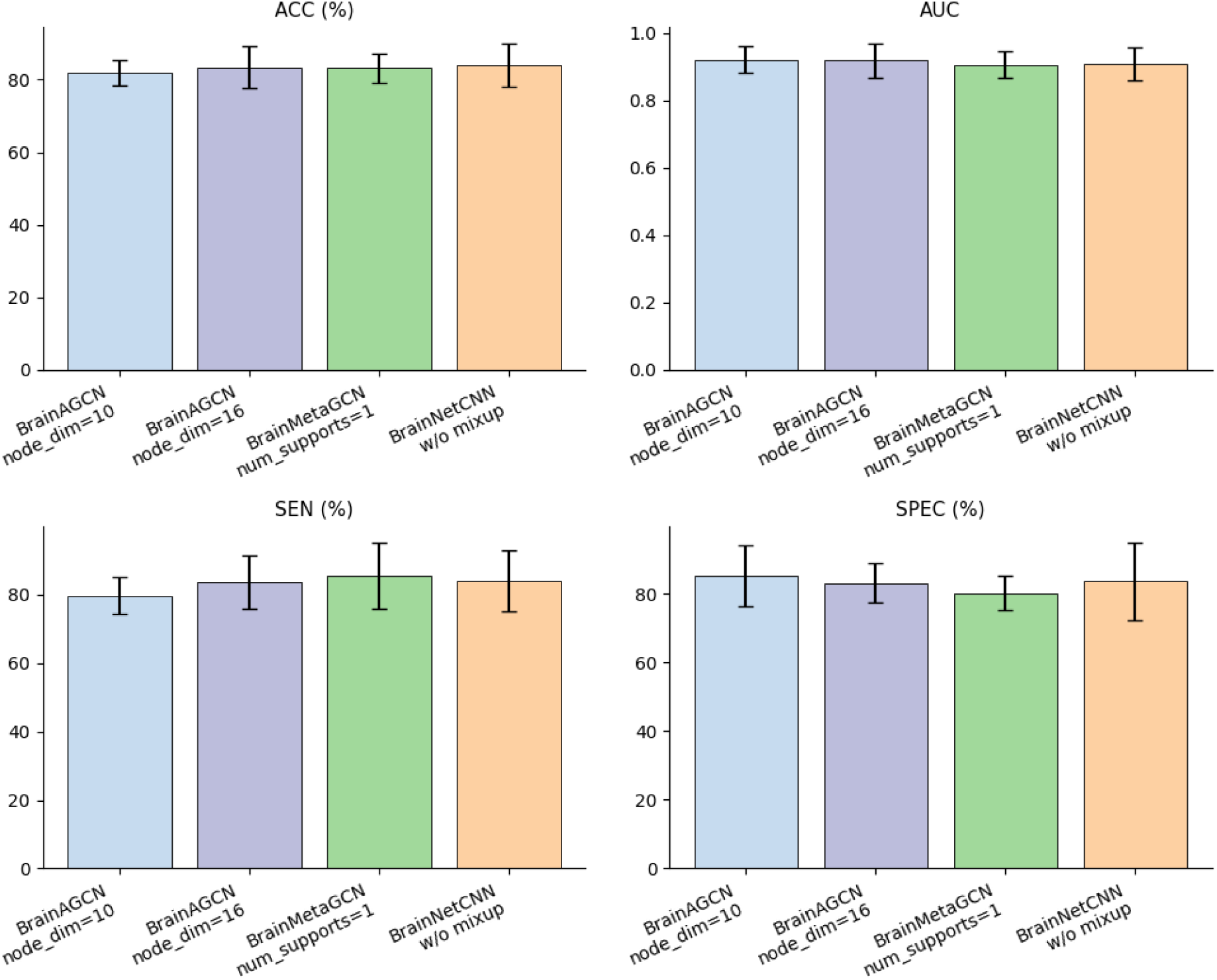
Ablation and hyperparameter sensitivity analysis. Ablation study evaluating the effects of key model components and hyperparameter settings on classification performance. Performance metrics (ACC, AUC, SEN, and SPEC) are compared across different node embedding dimensions, numbers of meta-graph supports, and with or without mixup regularization. Results demonstrate that the memory-augmented Meta-GCN configuration achieves superior and more stable performance, highlighting the importance of memory-based representation learning and lightweight architectural design for robust AMDD classification under small-sample conditions.

Removal of critical components, such as mixup regularization or memory augmentation, led to noticeable performance degradation, particularly in sensitivity and specificity. These results underscore the importance of memory-based representation learning and lightweight architectural design in mitigating noise and overfitting in rs-fMRI data. Overall, the ablation results confirm that the proposed configuration strikes a favorable balance between model complexity and generalization capability, making it well suited for AMDD classification under realistic small-sample conditions.

## Discussion

In the present study, we proposed a memory-augmented Meta-GCN framework for the classification of adolescent major depressive disorder (AMDD) using resting-state functional connectivity. By integrating dynamic meta-graph construction with a memory neural network mechanism, the proposed approach effectively addressed two central challenges in adolescent neuroimaging research: pronounced inter-individual heterogeneity and limited sample size. Across multiple independent datasets and evaluation settings, our model consistently outperformed state-of-the-art deep learning methods, while simultaneously yielding biologically interpretable network representations. These findings underscore the potential of lightweight, memory-enhanced graph models as a robust computational strategy for individualized neuropsychiatric diagnosis in adolescence.

### Methodological implications: individualized yet generalizable representation learning

A key methodological contribution of this work lies in the introduction of memory-based population modeling into functional brain network analysis. Conventional graph convolutional approaches typically assume a fixed or weakly adaptive graph structure and rely on large datasets to learn stable representations (*12*, *13*). Such assumptions are often violated in AMDD, where neurodevelopmental trajectories and symptom profiles vary substantially across individuals (*14*). By contrast, the memory module in our framework compresses population-level connectivity patterns into a set of prototypical templates and represents each individual as a weighted combination of these templates (*13*). This design enables the model to retain subject-specific characteristics while anchoring representations to stable population priors, thereby enhancing generalizability.

Importantly, the lightweight architecture and low-rank factorization employed in the proposed framework reduce model complexity without sacrificing discriminative power (*15*). The ablation analyses further demonstrate that both memory augmentation and dynamic meta-graph construction are essential for achieving robust performance, particularly in sensitivity and specificity. These results suggest that incorporating structured inductive biases—rather than increasing model depth or parameter count—may be a more effective strategy for rs-fMRI–based psychiatric classification under realistic small-sample conditions.

### Neurobiological interpretability and alignment with cortical functional hierarchy

Beyond classification performance, the proposed framework yields network representations that align with established principles of large-scale brain organization. We observed a significant association between memory-module–derived network weights and cortical functional hierarchy, indicating that the learned connectivity templates capture biologically meaningful gradients from unimodal sensory regions to transmodal association cortex (*16*). This finding is particularly relevant in the context of adolescence, a developmental period marked by ongoing maturation of higher-order association networks, including the default mode and executive control systems. Disruptions along the cortical hierarchy have been increasingly implicated in mood disorders, where aberrant integration between lower-level perceptual systems and higher-order cognitive–affective networks may underlie impairments in emotion regulation and self-referential processing (*6*). The alignment between our model-derived representations and functional hierarchy suggests that the memory-augmented Meta-GCN captures core neurodevelopmental features relevant to AMDD, rather than relying on spurious or dataset-specific patterns.

### Biological processes implicated by discriminative network components

Functional enrichment analysis of the discriminative network components further supports the biological plausibility of our findings. Enriched pathways related to synaptic transmission, axon guidance, and receptor tyrosine kinase signaling point to alterations in neural connectivity and plasticity, consistent with prior evidence of disrupted synaptic maturation and circuit refinement in adolescent depression (*17*). The involvement of immune- and cytokine-related signaling pathways aligns with emerging models that emphasize neuroimmune interactions in the pathophysiology of depression, particularly during sensitive developmental windows (*18*). Notably, these enriched processes span both neurodevelopmental and neuromodulatory domains, suggesting that AMDD may reflect the convergence of delayed or dysregulated brain maturation and altered molecular signaling (*19*). The ability of the proposed framework to bridge macroscale functional networks with microscale biological processes highlights its potential utility for integrative, multilevel investigations of psychiatric disorders.

### Clinical and translational relevance

From a clinical perspective, the proposed framework offers several advantages over existing neuroimaging-based diagnostic approaches. First, its improved sensitivity suggests enhanced capability to identify adolescents with depression, which is critical given the high rates of underdiagnosis and misdiagnosis in this population (*2*). Second, the interpretability of the learned network representations facilitates mechanistic insight, a prerequisite for clinical trust and translational adoption (*20*). Rather than functioning as a “black box,” the model yields anatomically and biologically meaningful features that can be related to established neurodevelopmental theories of depression (*6*, *9*). Moreover, the individualized representation enabled by the memory module may prove valuable for precision psychiatry applications, such as symptom subtyping, prognosis estimation, or treatment response prediction. By capturing both shared and individual-specific connectivity patterns, the framework lays the groundwork for future studies aiming to move beyond binary diagnosis toward personalized characterization of adolescent depression.

### Limitations and future directions

Several limitations should be acknowledged. First, although multiple independent datasets were used to assess generalizability, all participants were of Han Chinese ethnicity, which may limit the applicability of the findings to other populations. Future studies should validate the proposed framework in more diverse, multi-site cohorts. Second, the cross-sectional design precludes direct inference regarding developmental trajectories or illness progression. Longitudinal data would be essential to fully exploit the memory module’s capacity to encode temporal and stage-dependent patterns. Additionally, while functional enrichment analyses provide valuable biological context, they rely on indirect mapping between functional networks and molecular processes. Integrating multimodal data, such as structural connectivity, transcriptomic profiles, or neurochemical imaging, may further enhance mechanistic interpretation. Finally, although the model is lightweight relative to many deep learning approaches, real-world clinical deployment will require further optimization, external validation, and prospective testing.

## Conclusions

In summary, this study introduces a memory-augmented Meta-GCN framework that achieves robust, interpretable, and generalizable classification of adolescent major depressive disorder using resting-state fMRI data. By jointly addressing inter-individual heterogeneity and small-sample constraints, the proposed approach advances both methodological and neurobiological understanding of AMDD. These findings highlight the promise of memory-based graph learning as a powerful tool for precision neuropsychiatry and open new avenues for investigating the developmental mechanisms underlying adolescent depression.

## Methods

### Exploratory Dataset 1

#### Exploratory Dataset 1

The discovery sample was assembled at Renmin Hospital of Wuhan University and included adolescents aged 11–17 years. Patients in the depression group met diagnostic criteria for major depressive disorder at their first episode and had not previously received antidepressant treatment. Diagnosis was established through structured psychiatric assessment and confirmed by certified psychiatrists. Only participants with clinically significant depressive symptoms were retained for analysis. Healthy controls were recruited from the same catchment area and were frequency-matched to the patient group on major demographic factors. For both groups, exclusion criteria covered major neurological disease, severe medical illness, substance misuse, MRI contraindications, and other psychiatric disorders that could confound interpretation of the imaging findings. After exclusion of scans with excessive motion and records with incomplete key variables, the final discovery cohort comprised 302 adolescents with MDD (mean age = 15.40 ± 2.09 years; 134 females) and 207 healthy controls (HC, mean age = 15.33 ± 1.77 years; 86 females). Additional recruitment details and exclusion procedures are provided in the Supplementary Materials.

#### Replication Dataset 2

To assess reproducibility, we analyzed an external cohort from the NIMH Characterization and Treatment of Adolescent Depression study available through OpenNeuro (ds004627)(*21*). From this resource, we selected participants classified as having MDD or as healthy controls after applying the same quality-control principles used in the discovery dataset. Following motion screening and sample curation, 73 adolescents with MDD (mean age = 16.80 ± 1.22 years; 31 females) and 28 healthy controls (HC; mean age = 16.82 ± 1.40 years; 12 females) were included in the replication analyses. Further information regarding the public dataset and preprocessing can be found in the Supplementary Materials and the original dataset documentation (*21*).

#### Data preprocessing, quality control, and functional connectome

Imaging data were organized in BIDS (*22*) format and processed with a standardized pipeline based on fMRIPrep (*23*). Structural preprocessing included bias-field correction, skull stripping, tissue segmentation, surface reconstruction, and normalization to template space (*24*). Functional images underwent slice-timing correction, motion correction, and registration to each participant’s anatomical image (*25*). For connectome analysis, denoised resting-state signals were extracted from the 400-region Schaefer cortical atlas. Nuisance regression and temporal filtering were performed using Nilearn, after which pairwise Pearson correlations were calculated across all regional time series. Correlation matrices were then transformed to z values and used as subject-level functional connectivity representations for subsequent modeling. Detailed procedures for data quality control are provided in the Supplementary Materials (Data Quality Control section).

#### Functional Brain Network Construction

Resting-state functional magnetic resonance imaging (rs-fMRI) data were used to construct functional brain networks. The whole brain was parcellated into N regions of interest (ROIs) based on a predefined brain atlas. For each ROI, blood oxygen level-dependent (BOLD) time series were extracted, and Pearson correlation coefficients were calculated between every pair of ROIs to quantify functional connectivity. For two time series x and y of length T, the Pearson correlation coefficient is defined as:

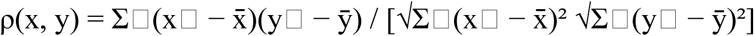

where x^-^ and ȳ denote the mean values of x and y, respectively. The resulting functional connectivity matrix FCN ∈ ℝᴺ×ᴺ was further normalized using Fisher’s r-to-z transformation. The functional brain network was modeled as a graph G = (V, E, A), where V represents the set of ROIs, E denotes the edges encoding functional connections, and A ∈ ℝᴺ×ᴺ is a learnable weighted adjacency matrix. Node features were defined as row vectors of the FCN, forming the node feature matrix X ∈ ℝᴺ×ᴺ.

#### Overall Framework of MAMGL

We propose a Memory-Augmented Meta-Graph Learning (MAMGL) framework for adolescent major depressive disorder diagnosis. The framework consists of three core components: a Meta-Graph Generator, a Meta-Graph Convolutional Network (Meta-GCN), and a Memory Augmentation Module (*11*). By introducing a learnable meta-node memory pool, the proposed framework effectively captures both population-level commonalities and individual-specific characteristics under limited sample conditions.

#### Meta-Graph Generator

To avoid biases introduced by predefined or heuristic graph sparsification strategies, a learnable meta-graph generation mechanism was developed. A meta-node memory pool Φ ∈ ℝᵐ×ᴰΦ (m ≪ N) was first constructed, where m denotes the number of memory items and DΦ represents the embedding dimension.

Two hyper-networks were employed to generate source and target node embeddings:

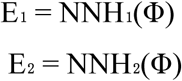

where NNH₁ and NNH₂ are fully connected hyper-networks parameterized by W₁ and W₂ ∈ ℝᴺ×ᵐ, respectively.

To model directed graph structures, two normalized meta-graphs were computed:

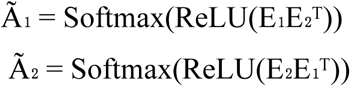

The resulting adjacency matrix Ã ∈ ℝᴺ×ᴺ represents adaptive weighted connections between brain regions.

#### Meta-Graph Convolutional Network

Based on the generated meta-graph, node representations were learned using a Meta-GCN. A single meta-graph convolutional layer is defined as:

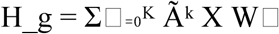

where K denotes the propagation depth and W1 are learnable parameters. Bidirectional graph convolution was adopted to capture directed dependencies:

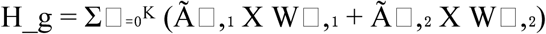

The Meta-GCN consists of two stacked meta-graph convolutional layers followed by a residual connection:

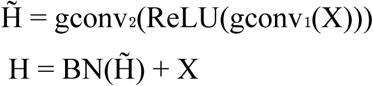

where BN denotes batch normalization.

#### Memory Augmentation Module

To further enhance discriminative node representations, a memory augmentation mechanism was introduced. First, the Meta-GCN output was projected to a lower-dimensional space:

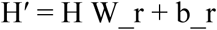

Then, query vectors were obtained:

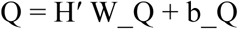

Attention weights between queries and the memory pool were computed as:

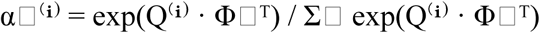

The memory representation was obtained by weighted summation:

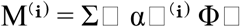

The memory-enhanced representation was formed by concatenation:

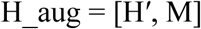

#### Feature Fusion and Classification

A two-layer multilayer perceptron (MLP) was applied to fuse the concatenated features, producing the final node-level representation. The fused features were flattened and fed into a three-layer MLP classifier to output binary classification results (major depressive disorder or healthy control).

#### Function gradient definition

Macroscale cortical gradients were estimated with BrainSpace(v0.1.10) (*26*). After z transformation of the connectivity matrices, each regional connectivity profile was sparsified by retaining its strongest connections (*27*). Inter-regional similarity was then quantified with a normalized-angle affinity kernel, and low-dimensional gradient components were obtained using diffusion map embedding (*28*). To enable group-level comparison, participant-specific gradients were aligned to a template derived from the mean connectome.

#### Enrichment analysis of network pattern of meta-GCN

Cell-type fractions were estimated by deconvolving microarray samples obtained from the Allen Human Brain Atlas (AHBA; http://human.brain-map.org/) (*29*). To investigate how functional reorganization is genetically regulated, AHBA transcriptomic data were integrated with high-resolution brain connectivity patterns. Regional microarray expression data were derived from six postmortem human brains (mean age = 42.50 ± 13.38 years; male/female = 5/1), comprising 3,702 spatially distinct tissue samples. Data preprocessing and mapping were performed using the abagen toolbox (*30*) (https://github.com/netneurolab/abagen), and gene expression profiles were mapped onto 400 cortical regions defined by the Schaefer 400-parcellation scheme. Subsequently, network patterns derived from the Meta-GCN framework were correlated with cortical gene expression profiles.

We next examined whether genes exhibiting transcriptional dysregulation at the messenger RNA (mRNA) level in postmortem brain tissue were preferentially expressed in cortical regions associated with gradient abnormalities across different subtypes. Functional pathway enrichment analysis was conducted using Metascape (https://metascape.org/gp/index.html#/main/step1), an automated meta-analysis platform integrating over 40 independent knowledge bases to identify shared or subtype-specific biological pathways (*31*). Genes most strongly associated with functional reorganization patterns were submitted to Metascape, and enriched pathways were thresholded at a false discovery rate (FDR)–corrected significance level of 5%, followed by validation using a null model.

This study further aimed to delineate the topographic correlations between Meta-GCN–derived network patterns and other salient neural features. To ensure robust statistical inference, we implemented a spatial null model designed to systematically disrupt the correspondence between two topographic maps while preserving their intrinsic spatial autocorrelation (*32*). Specifically, maps of key neural features were randomly permuted and their relationships with network patterns were recalculated. These spatial coordinates served as the basis for generating null distributions via randomly sampled rotations, with node values reassigned to the nearest parcels. This procedure was repeated 1,000 times to ensure stability. Notably, rotational transformations were first applied to one hemisphere and then mirrored onto the contralateral hemisphere. The 95th percentile of occurrence frequencies derived from both spatial and temporal null models was used as the threshold for statistical significance. This stringent thresholding strategy enabled the identification of meaningful topographic correlations while adequately controlling for inherent spatial autocorrelation in the data.

## Data and Code Availability

Because the clinical imaging data contain sensitive participant information, access to the non-public cohort is available from the corresponding author upon reasonable request and subject to institutional approval. The external replication dataset is publicly available from OpenNeuro (ds004627). Public reference resources used in this study, including AHBA, BrainSpace, neuromaps, and the ENIGMA toolbox, can be obtained from their respective project websites. Code supporting the analyses is available at the project repository.

## Acknowledgments

Xiaobo Liu is supported by the China Scholarship Council. This work was supported in part by the Health of Hubei Province Scientific Research Project under Grant 2020Cfb512, and by the Mental Health Research Institute of Three Gorges University: YCXL-23-11. We thank the National Center for Protein Sciences at Peking University in Beijing, China for assistance on data analysis.

## Exploratory Dataset 1

### Exploratory Dataset 1

Participants were recruited through the Center for the Prevention and Management of Depression at Renmin Hospital of Wuhan University using a combination of public advertisements and clinical referrals. Data collection was conducted between May 4, 2018, and December 30, 2023. The initial sample comprised 361 adolescents diagnosed with major depressive disorder (MDD) and 252 healthy controls (HC). MDD diagnoses were established according to DSM-IV criteria using the Structured Clinical Interview for DSM-IV Axis I Disorders (SCID) and were independently confirmed by two board-certified psychiatrists. In addition, all MDD participants were assessed by two independent board-certified child and adolescent psychiatrists using the Mini International Neuropsychiatric Interview for Children and Adolescents (MINI-KID, Chinese version), based on DSM-5 diagnostic criteria.

Eligible participants were between 11 and 17 years of age, had a duration of depressive symptoms of less than 12 months, and had no prior history of antidepressant medication or electroconvulsive therapy. Exclusion criteria included comorbid psychiatric disorders, history of head trauma with loss of consciousness, somatic illnesses affecting brain morphology, substance abuse, left-handedness, severe physical or neurological disorders, contraindications to magnetic resonance imaging (MRI), and recent medication use within the preceding five half-lives. Depression severity was assessed using the 17-item Hamilton Rating Scale for Depression (HAMD-17), and a minimum score of 17 was required for inclusion in the MDD group.

For the MDD cohort, 45 participants were excluded due to excessive head motion during resting-state functional magnetic resonance imaging (rs-fMRI) acquisition (mean framewise displacement > 2 mm). An additional 14 participants were excluded because of missing essential demographic or clinical information (e.g., age, sex, or HAMD scores). The final exploratory dataset therefore consisted of 302 adolescents with MDD. Following identical quality control procedures, 207 healthy controls matched for age, sex, ethnicity, educational level, and handedness were randomly recruited from the local community and included in the final analysis.

### Imaging Acquisition

Resting-state fMRI data were acquired at the PET Center of Renmin Hospital of Wuhan University using a 3.0-T General Electric MRI scanner, following previously established imaging protocols. During scanning, participants were instructed to lie supine with their eyes closed, remain awake, and minimize head movement. Functional images were obtained using a gradient-echo echo-planar imaging (EPI) sequence with the following parameters: repetition time (TR) = 2000 ms; echo time (TE) = 30 ms; flip angle = 90°; field of view = 240 × 240 mm²; matrix size = 64 × 64; 32 axial slices; slice thickness = 3.0 mm with no inter-slice gap. A total of 212 volumes were acquired over approximately 16 minutes.

### Replication Dataset 2

To validate the findings from the exploratory dataset, we analyzed an independent replication sample derived from the Characterization and Treatment of Adolescent Depression (CAT-D) study, accessed via OpenNeuro (dataset ID: ds004627). The CAT-D study is an ongoing longitudinal investigation recruiting adolescents through community practitioner referrals, self-referrals, and advertisements primarily across Maryland, Virginia, and the District of Columbia.

Participants were eligible if they were between 11 and 17 years of age at enrollment and met criteria for current or past MDD, subthreshold MDD (s-MDD), or were healthy controls. As of March 11, 2021, a total of 279 participants had been enrolled. For the present analyses, we included adolescents classified as HC or with a current or lifetime diagnosis of MDD based on clinician-administered Kiddie Schedule for Affective Disorders and Schizophrenia–Present and Lifetime Version (KSADS-PL) assessments.

To examine data collected before and during the COVID-19 pandemic, only participants with available assessments in both periods were included. The World Health Organization’s declaration of COVID-19 as a global pandemic on March 11, 2020, was used as the temporal reference. Pre-pandemic assessments were defined as those conducted between March 11, 2019, and March 10, 2020, while pandemic assessments were defined as those conducted between March 11, 2020, and March 11, 2021. Based on these criteria, 166 participants were initially identified for analyses examining associations between the pandemic and depressive and anxiety symptoms.

### Data Quality Control

All rs-fMRI data underwent identical quality control procedures. Participants with excessive head motion, defined as a mean framewise displacement greater than 2 mm, were excluded. In the exploratory dataset, 44 adolescents with MDD and 45 healthy controls were excluded on this basis, resulting in a final sample of 302 MDD participants and 207 HC. In the replication dataset, 16 participants were excluded due to excessive head motion, yielding a final sample of 101 adolescents, including 73 with MDD and 28 healthy controls.

---

**Supplementary Table 1.** Demographic information for datasets used in this study. (HAMD, Hamilton Depression Rating Scale; HAMA, Hamilton Anxiety Rating Scale)

|  | Group | Number | Age | Sex (F/M) | HAMD | HAMA |
| --- | --- | --- | --- | --- | --- | --- |
| Exploration dataset | A-M DD | 302 | 15.40±2.09 | 134/156 | 25.11±5.63 | 23.23±6.50 |
|  | HC | 207 | 15.53±1.98 | 82/121 | ---- | ---- |
| | P values | | $p > 0.05$ | $p > 0.05$ | ---- | ---- |
| Replication dataset | A-M DD | 73 | 16.80±1.22 | 31/42 | ---- | ---- |
|  | HC | 28 | 16.82±1.40 | 12/16 | ---- | ---- |
| | P values | | $p > 0.05$ | $p > 0.05$ | ---- | ---- |

## Reference

1. D. M. Hafeman, J. Merranko, T. R. Goldstein, D. Axelson, B. I. Goldstein, K. Monk, M. B. Hickey, D. Sakolsky, R. Diler, S. Iyengar, D. A. Brent, D. J. Kupfer, M. W. Kattan, B. Birmaher, Assessment of a Person-Level Risk Calculator to Predict New-Onset Bipolar Spectrum Disorder in Youth at Familial Risk. JAMA Psychiatry 74, 841–847 (2017).

2. A. Thapar, S. Collishaw, D. S. Pine, A. K. Thapar, Depression in adolescence. The Lancet 379, 1056–1067 (2012).

3. S. P. Ahmed, A. Bittencourt-Hewitt, C. L. Sebastian, Neurocognitive bases of emotion regulation development in adolescence. Dev. Cogn. Neurosci. 15, 11–25 (2015).

4. A.-K. Allgaier, B. Frühe, K. Pietsch, B. Saravo, M. Baethmann, G. Schulte-Körne, Is the Children’s Depression Inventory Short version a valid screening tool in pediatric care? A comparison to its full-length version. J. Psychosom. Res. 73, 369–374 (2012).

5. C. H. Miller, J. P. Hamilton, M. D. Sacchet, I. H. Gotlib, Meta-analysis of Functional Neuroimaging of Major Depressive Disorder in Youth. JAMA Psychiatry 72, 1045–1053 (2015).

6. X. Liu, B. Wan, X. Wu, X. Zhang, L. Liu, S. Long, R. Ge, R. Cui, X. Wen, X. Liu, W. Peng, G. Yang, Y. Gao, Subtypes of adolescent major depressive disorder characterized by divergent information dynamics in sensory-association cortices. Nat. Commun., doi: 10.1038/s41467-026-69697-2 (2026).

7. X. Liu, B. Wan, X.-H. Zhang, R. Cui, S. Long, R. Ge, L. Liu, J. Xiao, Z.-Q. Liu, J. Yan, K. Xie, M. Yao, X. Wen, S. Wang, Y. Gao, Episode-specific cortical functional connectome reorganization and neurobiological correlates in bipolar disorder: a cross-sectional study. BMC Med. 23, 457 (2025).

8. K. Qin, D. Lei, W. H. L. Pinaya, N. Pan, W. Li, Z. Zhu, J. A. Sweeney, A. Mechelli, Q. Gong, Using graph convolutional network to characterize individuals with major depressive disorder across multiple imaging sites. eBioMedicine 78, 103977 (2022).

9. X. Liu, Y. Xu, H. Xu, L. He, S. Long, Y. Huang, Y. Wang, Y. Lu, Y. Huang, J. Wu, H. Gao, X. Liu, Advancing interpretable cardiac disease diagnosis via a transformer-convolutional hybrid network on electrocardiograms. Eng. Appl. Artif. Intell. 152, 110675 (2025).

10. S. Sukhbaatar, A. Szlam, J. Weston, R. Fergus, End-To-End Memory Networks. arXiv arXiv:1503.08895 [Preprint] (2015). 10.48550/arXiv.1503.08895.

11. J. Weston, S. Chopra, A. Bordes, Memory Networks. arXiv arXiv:1410.3916 [Preprint] (2015). 10.48550/arXiv.1410.3916.

12. D. Zhi, V. D. Calhoun, C. Wang, X. Li, X. Ma, L. Lv, W. Yan, D. Yao, S. Qi, R. Jiang, J. Zhao, X. Yang, Z. Lin, Y. Zhang, Y. C. Chung, C. Zhuo, J. Sui, “BNCPL: Brain-Network-based Convolutional Prototype Learning for Discriminating Depressive Disorders” in 2021 43rd Annual International Conference of the IEEE Engineering in Medicine & Biology Society (EMBC) (2021; https://ieeexplore.ieee.org/document/9630010), pp. 1622–1626.

13. K. Zheng, S. Yu, L. Chen, L. Dang, B. Chen, BPI-GNN: Interpretable brain network-based psychiatric diagnosis and subtyping. NeuroImage 292, 120594 (2024).

14. S. P. Ahmed, A. Bittencourt-Hewitt, C. L. Sebastian, Neurocognitive bases of emotion regulation development in adolescence. Dev. Cogn. Neurosci. 15, 11–25 (2015).

15. W. Tu, F. Fu, L. Kong, B. Jiang, D. Cobzas, C. Huang, Low-Rank Plus Sparse Decomposition of fMRI Data With Application to Alzheimer’s Disease. Front. Neurosci. 16 (2022).

16. D. S. Margulies, S. S. Ghosh, A. Goulas, M. Falkiewicz, J. M. Huntenburg, G. Langs, G. Bezgin, S. B. Eickhoff, F. X. Castellanos, M. Petrides, E. Jefferies, J. Smallwood, Situating the default-mode network along a principal gradient of macroscale cortical organization. Proc. Natl. Acad. Sci. 113, 12574–12579 (2016).

17. J. M. Loftis, M. Huckans, B. J. Morasco, Neuroimmune mechanisms of cytokine-induced depression: Current theories and novel treatment strategies. Neurobiol. Dis. 37, 519–533 (2010).

18. R. Nusslock, L. B. Alloy, G. H. Brody, G. E. Miller, Annual Research Review: Neuroimmune network model of depression: a developmental perspective. J. Child Psychol. Psychiatry 65, 538–567 (2024).

19. H. C. Brenhouse, J. M. Schwarz, Immunoadolescence: Neuroimmune development and adolescent behavior. Neurosci. Biobehav. Rev. 70, 288–299 (2016).

20. Y. Xiao, F. Y. Womer, S. Dong, R. Zhu, R. Zhang, J. Yang, L. Zhang, J. Liu, W. Zhang, Z. Liu, X. Zhang, F. Wang, A neuroimaging-based precision medicine framework for depression. *Asian J*. Psychiatry 91, 103803 (2024).

21. N. Sadeghi, P. Q. Fors, L. Eisner, J. Taigman, K. Qi, L. S. Gorham, C. C. Camp, G. O’Callaghan, D. Rodriguez, J. McGuire, E. M. Garth, C. Engel, M. Davis, K. E. Towbin, A. Stringaris, D. M. Nielson, Mood and Behaviors of Adolescents With Depression in a Longitudinal Study Before and During the COVID-19 Pandemic. J. Am. Acad. Child Adolesc. Psychiatry 61, 1341–1350 (2022).

22. K. J. Gorgolewski, T. Auer, V. D. Calhoun, R. C. Craddock, S. Das, E. P. Duff, G. Flandin, S. S. Ghosh, T. Glatard, Y. O. Halchenko, D. A. Handwerker, M. Hanke, D. Keator, X. Li, Z. Michael, C. Maumet, B. N. Nichols, T. E. Nichols, J. Pellman, J.-B. Poline, A. Rokem, G. Schaefer, V. Sochat, W. Triplett, J. A. Turner, G. Varoquaux, R. A. Poldrack, The brain imaging data structure, a format for organizing and describing outputs of neuroimaging experiments. Sci. Data 3, 160044 (2016).

23. O. Esteban, C. J. Markiewicz, R. W. Blair, C. A. Moodie, A. I. Isik, A. Erramuzpe, J. D. Kent, M. Goncalves, E. DuPre, M. Snyder, H. Oya, S. S. Ghosh, J. Wright, J. Durnez, R. A. Poldrack, K. J. Gorgolewski, fMRIPrep: a robust preprocessing pipeline for functional MRI. Nat. Methods 16, 111–116 (2019).

24. K. Gorgolewski, C. D. Burns, C. Madison, D. Clark, Y. O. Halchenko, M. L. Waskom, S. S. Ghosh, Nipype: A Flexible, Lightweight and Extensible Neuroimaging Data Processing Framework in Python. *Front*. Neuroinformatics 5 (2011).

25. H.-T. Wang, S. L. Meisler, H. Sharmarke, N. Clarke, N. Gensollen, C. J. Markiewicz, F. Paugam, B. Thirion, P. Bellec, Continuous evaluation of denoising strategies in resting-state fMRI connectivity using fMRIPrep and Nilearn. PLOS Comput. Biol. 20, e1011942 (2024).

26. R. Vos de Wael, O. Benkarim, C. Paquola, S. Lariviere, J. Royer, S. Tavakol, T. Xu, S.-J. Hong, G. Langs, S. Valk, B. Misic, M. Milham, D. Margulies, J. Smallwood, B. C. Bernhardt, BrainSpace: a toolbox for the analysis of macroscale gradients in neuroimaging and connectomics datasets. *Commun*. Biol. 3, 103 (2020).

27. D. S. Margulies, S. S. Ghosh, A. Goulas, M. Falkiewicz, J. M. Huntenburg, G. Langs, G. Bezgin, S. B. Eickhoff, F. X. Castellanos, M. Petrides, E. Jefferies, J. Smallwood, Situating the default-mode network along a principal gradient of macroscale cortical organization. Proc. Natl. Acad. Sci. 113, 12574–12579 (2016).

28. R. R. Coifman, S. Lafon, Diffusion maps. Appl. Comput. Harmon. Anal. 21, 5–30 (2006).

29. E. H. Shen, C. C. Overly, A. R. Jones, The Allen Human Brain Atlas: Comprehensive gene expression mapping of the human brain. Trends Neurosci. 35, 711–714 (2012).

30. R. D. Markello, A. Arnatkeviciute, J.-B. Poline, B. D. Fulcher, A. Fornito, B. Misic, Standardizing workflows in imaging transcriptomics with the abagen toolbox. eLife 10, e72129 (2021).

31. Y. Zhou, B. Zhou, L. Pache, M. Chang, A. H. Khodabakhshi, O. Tanaseichuk, C. Benner, S. K. Chanda, Metascape provides a biologist-oriented resource for the analysis of systems-level datasets. Nat. Commun. 10, 1523 (2019).

32. J. Y. Hansen, G. Shafiei, R. D. Markello, K. Smart, S. M. L. Cox, M. Nørgaard, V. Beliveau, Y. Wu, J.-D. Gallezot, É. Aumont, S. Servaes, S. G. Scala, J. M. DuBois, G. Wainstein, G. Bezgin, T. Funck, T. W. Schmitz, R. N. Spreng, M. Galovic, M. J. Koepp, J. S. Duncan, J. P. Coles, T. D. Fryer, F. I. Aigbirhio, C. J. McGinnity, A. Hammers, J.-P. Soucy, S. Baillet, S. Guimond, J. Hietala, M.-A. Bedard, M. Leyton, E. Kobayashi, P. Rosa-Neto, M. Ganz, G. M. Knudsen, N. Palomero-Gallagher, J. M. Shine, R. E. Carson, L. Tuominen, A. Dagher, B. Misic, Mapping neurotransmitter systems to the structural and functional organization of the human neocortex. Nat. Neurosci. 25, 1569–1581 (2022).

